# Systems Biology Understanding of the Effects of Lithium on Cancer

**DOI:** 10.1101/363077

**Authors:** Weihao Ge, Eric Jakobsson

## Abstract

Lithium has many widely varying biochemical and phenomenological effects, suggesting that a systems biology approach is required to understand its action. Multiple lines of evidence point to lithium as a significant factor in development of cancer, showing that understanding lithium action is of high importance. In this paper we undertake first steps towards a systems approach by analyzing mutual enrichment between the interactomes of lithium-sensitive enzymes and the pathways associated with cancer. This work integrates information from two important databases, STRING and KEGG pathways. We find that for the majority of cancer pathways the mutual enrichment is many times greater than chance, reinforcing previous lines of evidence that lithium is an important influence on cancer.

## Introduction

### Clinical and Epidemiological Context for Lithium and Cancer

By far the most common medical use of lithium is as a first line therapy for bipolar disorder, including associated depression as well as mania.^1^ A comprehensive review of the literature confirms that lithium is also effective against unipolar depression with unique anti-suicidal effectiveness, and may also be useful against cancer and neurodegenerative disease.^2^

One line of evidence for the possible use of lithium as an anticancer agent is epidemiological. A retrospective study showing that psychiatric patients undergoing lithium therapy for bipolar disorder had a much lower incidence of cancer than a matched group not receiving lithium therapy. ^3^ More recent studies of similar design, one conducted nationwide across Sweden, and another across Taiwan, achieved the same result.^4 5^ On the other hand another nationwide study, this time from Denmark, showed no correlation of lithium with colorectal adenocarcinoma.^6^ On closer look, the Denmark study does not contradict the Swedish study. The Swedish study also found that for the entire population lithium was not correlated with cancer incidence, but in addition found that bipolar individuals not treated with lithium had a higher incidence of cancer than the general population. Lithium-treated bipolar patients, on the other hand, had essentially the same cancer incidence as the general population.

One piece of experimental evidence for lithium’s potential as a cancer therapeutic modality is that it was observed to inhibit prostate tumor growth,^7^ presumably through its ability to inhibit GSK3. A detailed study of molecular mechanisms by which lithium inhibition of GSK3-beta inhibits proliferation of prostate tumor cells in culture was presented by Sun et al.^8^ The work was subsequently extended to an animal model.^9^ A clinical trial for the effect of lithium coupled with prostatectomy on men has been conducted but as of this writing results have not yet been published.^10^

With respect to other cancers, lithium has been found to be lethal to neuroblastoma cells but not to normal nerve cells.^11^ The experimentally determined effective dose was 12 mM, a level which would be lethal if achieved systemically in a human or model organism but perhaps could be induced locally. A similar effect was found in ovarian cancer cells,^12^ although a subsequent similar study on ovarian cancer cells suggests only a more modest benefit.^13^ It is not clear from our reading of the two ovarian cancer papers why the results are significantly different from each other.

With respect to colorectal cancer, one study suggests that lithium inhibits proliferation of a colorectal cancer cell line.^14^ Another study on colon cancer cells showed that lithium specifically induced a reversal of the epithelial-to-mesenchymal transition characteristic of the cancer cells.^15^

Two studies with relatively small sample size suggested a possible link between lithium and tumors of the upper urinary tract.^16 17^ However a large-scale study involving all urinary tract cancers in Denmark over a multi-year period found no correlation with lithium use.^18^

Because lithium therapy is systemic rather than topical or local, it follows that lithium might inhibit metastasis. Evidence that this is the case for colon cancer comes from observation of inhibition of metastasis-inducing factors by lithium and by observation on reduced metastasis in model animals given lithium therapy.^19^

Autophagy is a key cellular process in the inhibition of cancer.^20^ Lithium has been shown to induce autophagy, due to its inhibition of inositol monophosphatase.^21^ The full range of lithium effects on autophagy is complicated,^22^ as might be expected because lithium has multiple targets, which themselves have multiple substrates.

Because of the promising indications as cited above, lithium has been suggested as one of a number of drugs commonly used for other reasons, to be repurposed for cancer.^23^

### Biochemical Context for Lithium and Cancer

Much of lithium’s known biochemical action may be summarized by noting that it inhibits some phosphate-transfer enzymes (primarily phosphatases and kinases) that have magnesium as a co-factor.^2^ A common underlying biophysical basis for competition between lithium and magnesium for modulating phosphate-transfer enzymes, is suggested by noting that the primary energy source for cells and the substrate for phosphorylating enzymes is not bare ATP, but rather magnesium-associated ATP (MgATP).^24^ NMR studies show that lithium associates with MgATP.^25^ Based on this admittedly small amount of data, we consider the possibility that lithium generally associates with magnesium-phosphate complexes and thus has the potential to modulate to some extent a large number of phosphorylation reactions and ATP-splitting processes.

Because mutations in G protein linked receptors have emerged as of interest in cancer research, ^26^ it is significant that lithium appears to inhibit β-adrenergic and muscarinic receptor coupling to G proteins by competing with magnesium, which facilitates such coupling. ^27 28 29 30 31^

In the literature we find evidence for direct lithium inhibition of seventeen human magnesium-dependent phosphate-transfer enzymes, as follows: A review by Phiel and Klein^32^ identified five (IMPase, IPPase, FBPase, BPntase, and GSK3B). Testing against a panel of 80 protein kinases^33^ revealed lithium sensitivity for eight more enzymes (MNK1, MNK2, smMLCK, PHK, CHK2, HIPK3, IKKϵ and TBK1). It has long been observed that adenyl cyclase activity is inhibited by lithium.^34^ Of nine different adenylyl cyclases tested, two (ADCY5 and ADCY7) are strongly inhibited by lithium and one (ADCY2) is less strongly but significantly inhibited.^35^ With the addition of GSK3A,^36^ we have a list of seventeen phosphate-transfer enzymes directly inhibited by lithium. An inspection of protein-protein interaction databases indicate that all seventeen interact with multiple other gene products. It should be noted that 72 out of 80 kinases^33^, and six out of nine adenylyl cyclases^35^, screened were found *not* to be lithium-sensitive.

Because lithium affects many different biological molecules and processes^2^, it is essential to utilize the tools of systems biology^37^ if a comprehensive understanding of lithium action and its prospects for therapy are to be obtained. Important concepts for organizing biological information in a systems context are pathways and networks. A very useful tool for obtaining data about known pathways is the KEGG database.^38^ An equally useful and complementary tool is the STRING database of interacting proteins.^39^

In the present paper we investigate further the possible linkages between lithium and cancer by analyzing the mutual enrichment between STRING-derived interactomes of lithium-sensitive enzymes, and the KEGG pathways associated with cancer.

## Methods

Analysis was performed on the interactomes of the above-mentioned lithium-sensitive genes. The interactomes of these genes were extracted from the STRING database (https://string-db.org). For each key gene, we adjust confidence level and order of neighbors (nearest only or next nearest included), so that each set contains a few hundred genes. This size is large enough for statistically reliable enrichment analysis.

## Disease Association

We used the R-package KEGGgraph^40 41^ to identify the genes associated with the cancer-relevant pathways.

### *P*-value calculation

The fundamental question we address is whether there is significant overlap or mutual enrichment between the interactomes of lithium-sensitive genes and the pathways or gene sets implicated in various cancers.

For each of the 17 lithium sets, an ensemble of 1000 null sets are generated by random selection from the human genome. Each null set is the same size as the corresponding lithium set. Then we used the R-package STRINGdb^42^ to perform KEGG pathway enrichment analysis. This operation is a particular example of the powerful technique of gene-annotation enrichment analysis. ^43^ In gene-annotation enrichment analysis a test list of genes (often derived from gene expression experiments) is compared to an organized database of gene annotations, often referred to as a gene ontology^44^, an array of gene lists corresponding to different biological functions, molecular functions, or locations in the cell. The output of the gene-annotation enrichment analysis is expressed as the likelihood that the list overlaps could have occurred by chance (p-value). A very low p-value implies that the degree of overlap is highly significant statistically and very likely is significant biologically. In our study the gene lists we are comparing are the interactomes of lithium sensitive enzymes on the one hand, and KEGG pathways associated with cancer on the other hand. For each KEGG term retrieved, a null distribution of uncorrected *p*-value is generated by the 1000 null sets. This gives us a measure of the false discovery rate, since any overlap between the null sets and the KEGG pathways is purely accidental. Then the fraction of null set uncorrected *p*-values smaller than or equal to the lithium-sensitive set uncorrected *p-*value would be the empirical *p*-value. For a detailed discussion of empirical p-value determination see Ge et al^45^.

## Results

Fig.1 shows mutual lithium interactome enrichment with specific cancer pathways, represented by heatmaps. Each area on the heatmap is a color-coded representation of the degree of mutual enrichment between the genes in the interactome of the indicated lithium sensitive enzyme and the genes in the indicated pathway. The darker the shade, the more significant the mutual enrichment of the interactome-pathway combination is. The light areas on the heatmap represent situations where a lithium-sensitive interactome has little or no mutual enrichment with a cancer pathway. The dark areas, deep orange and red, represent situations where enrichment is very strong—far greater than could be expected by chance.

It appears that the interactomes of five out of the 17 lithium-sensitive genes (ADCY2, ADCY5, ADCY7, BPNT1, and HIPK3) do not show significant mutual enrichment with specific cancer pathways. Chemical carcinogenesis shows significant mutual enrichment with only one of the interactomes, that of IMPAD1. For the remaining specific cancer pathways and lithium-sensitive interactomes, there are multiple areas of strong mutual enrichment. The genes contained in these overlapping areas, and their modes of regulation, appear worthy of further study in unraveling the details of the lithium vs cancer relationship.

In addition to the labeled specific cancer pathways we extended the analysis to signaling pathways in which dysfunction is implicated in cancer, as indicated in the literature.^46 47 48 49 50 51 52 53^ Figure 2 shows in heatmap form the mutual enrichment between the seventeen lithium-sensitive interactomes and thirteen pathways relevant to cancer.

**Figure 2.**
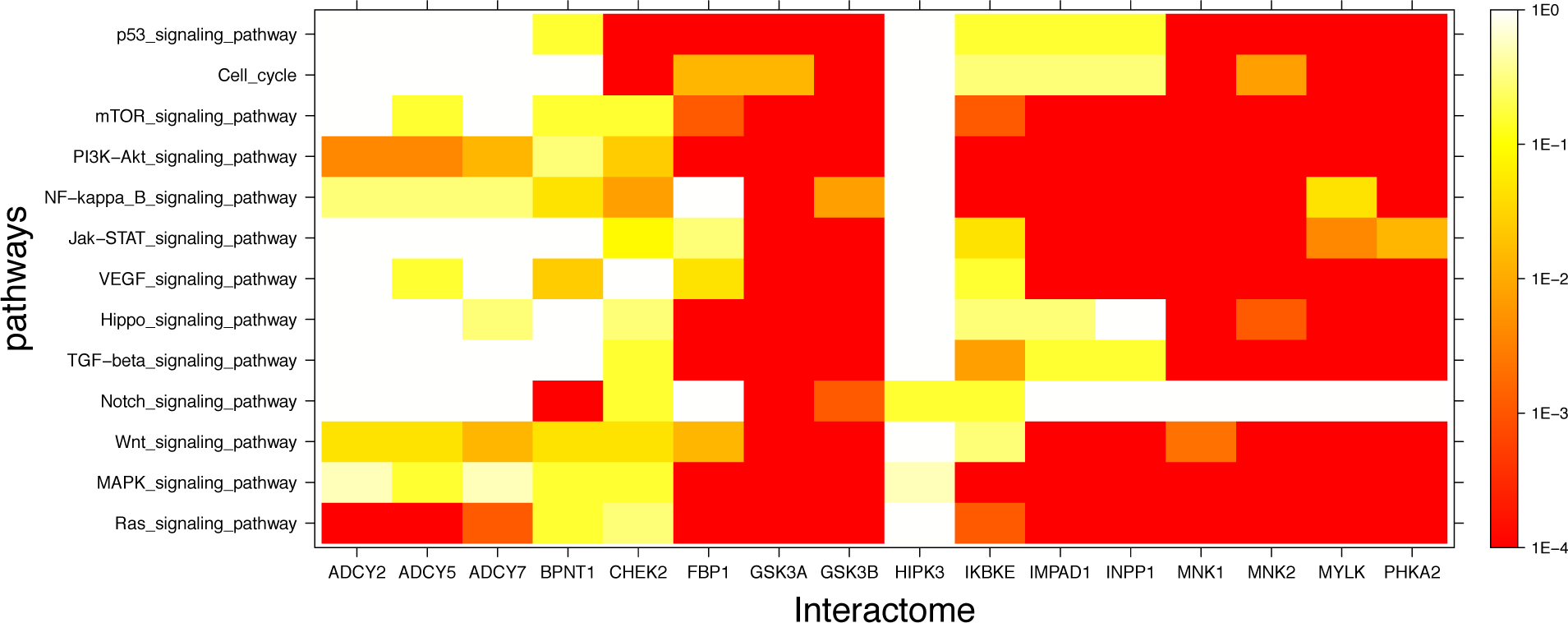
Visual representation of mutual enrichment patterns between signaling pathways implicated in cancer and the interactomes of lithium-sensitive gene products. Calibration of p-value vs. color is indicated by a vertical scale to the right of the heat map. Red or dark orange indicates very strong enrichment while lighter color indicates weak or, if white, no enrichment. Only one gene product appears not relevant to cancer, HIPK3. The three adenyl cyclases, BPNT1, and CHEK2 show strong mutual enrichment for only a couple of the pathways. Each of the remaining eleven interactomes show strong mutual enrichment with most of the cancer-relevant pathways.

The inescapable conclusion from Figures 1 and 2 is that variability in lithium concentration is likely to significantly modulate most cancer-relevant pathways. We should note that sensitivity to lithium does not necessarily imply a *beneficial* sensitivity. There are some indications for some cancers that lithium might be beneficial, as described in the Introduction section of this paper, but because of the complexity of the feedback relationships in these pathways, a complicated relationship between lithium ingestion and cancer incidence is very possible.

**Figure 1.**
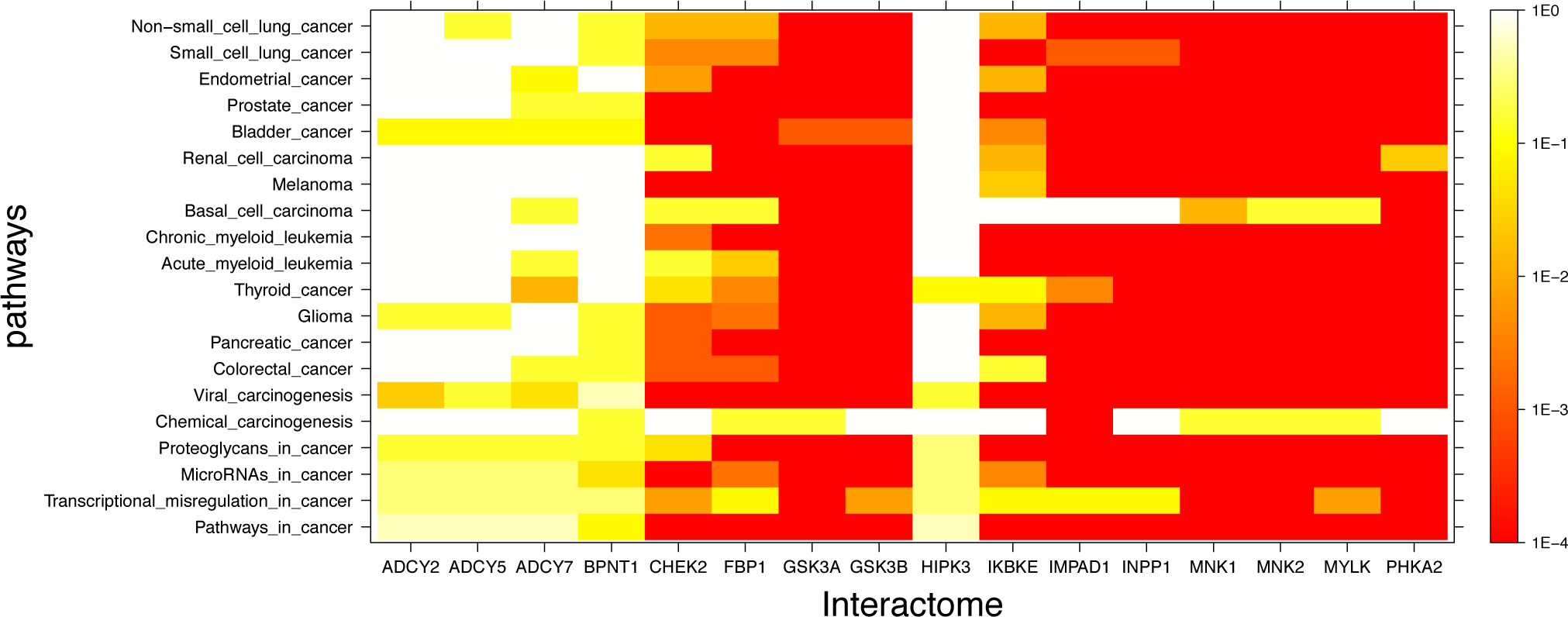
Visual representation of mutual enrichment patterns between specific cancer pathways and the interactomes of lithium-sensitive gene products. Calibration of p-value vs. color is indicated by a vertical scale to the right of the heat map. Red or dark orange indicates very strong enrichment while lighter color indicates weak or, if white, no enrichment. Five genes stand out as being not strongly connected to these cancer pathways: BPNT1, HIPK3, ADCY2, ADCY5, and ADCY7. Of the cancer pathways, chemical carcinogenesis stands out as being less likely to be strongly influenced by lithium levels, although there is a strong mutual enrichment between the interactome of IMPAD1 and this pathway. For the remainder of the genes and the remainder of the cancers, the relationship between the lithium-sensitive interactome and the cancer phenotype is strong.

## Summary and Discussion

We have conducted a pathway and network enrichment analysis exploring the role of lithium in multiple cancers and cancer-related pathways. The results show that for the large majority of such cancers, there is high mutual enrichment between the interactomes of lithium-sensitive enzymes and the pathways associated with those diseases, indicating that lithium is very likely to affect the incidence and course of the disease. Our results are consistent with a variety of lines of evidence from both epidemiology and from experiment, cited in the Introduction section of this paper, suggesting possible influence of lithium on the incidence and progression of cancer.

We hope that the results described in this paper will contribute to prioritizing and designing clinical trials of lithium for cancer. To provide context for such prioritization and design, it is essential to take into account the ways in which lithium is unique, both as a pharmaceutical and as an ion that is ubiquitous in the environment, and therefore ubiquitous in the water and food we ingest^2^:

1. Unlike other ions, lithium is not closely regulated by selective membrane transport processes. Rather it shares pathways that are mainly selective for other ions, in most cases sodium.^2^ Therefore, lithium concentration in both extracellular and intracellular compartments, rather than being nearly constant as is the case with other ions, is roughly proportional to lithium ingestion. ^54^ Whereas changes in the concentrations of other ions of more than a few percent have severe acute adverse consequences, the human body adjusts without acute adverse consequence to changes in lithium concentrations of several orders of magnitude. Our biochemistry has evolved to accommodate to widely varying lithium levels, as opposed to developing the ability to closely regulate lithium levels.
2. The multiple enzymes inhibited by lithium are each functionally linked to large numbers of other genes. This explains why the effects of lithium are widespread and varied; lithium has a modulating effect on many gene networks. We note that screening for lithium sensitivity has so far not included systematic examination of multiple variants of particular gene products, either mutational variants or alternative splices from the same gene. Therefore, it may be that some of the enzymes that have been found not lithium-sensitive may have mutational or splice variants that are sensitive. Conversely, some of the enzymes that have been found to be lithium-sensitive may have mutational or splice variants that are insensitive. The plausibility of such a possibility is exemplified by a functional, structural, and mutational study on an archaeal inositol monophosphatase.^55^ The archaeal enzyme has high homology (30% identical, 50% similar) to its human counterpart and functions in the same magnesium-dependent manner. In this study it was shown that a single amino acid substitution could convert the enzyme from its native lithium-insensitive form to a lithium-sensitive form. Perhaps of relevance, it has long been known that lithium responsiveness is significantly variable among human individuals.^56^
3. Unlike other pharmaceuticals, lithium is probably an essential trace element in the diet.^57 58 59^The question with lithium is not whether it should be ingested or not, but rather how much. Extreme lithium deprivation results in failure to thrive, while too much lithium is toxic. The existence of these extrema suggests existence of an intermediate optimum.

Therefore, we suggest that the correct question to ask with respect to lithium and a particular disease is not, “Should lithium be administered for this particular disease?” but rather, “What is the optimum blood level of lithium for this individual, given his or her disease history, status, genetic propensities, and other medications?” Unlike some pharmaceuticals that are more specific and inhibit or activate one gene or a small number of genes, the model for lithium action is that it alters the balance between a large number of interacting processes and pathways. Thus, a dose-response curve for lithium is likely to be highly nonlinear and not always monotonic.

There are just a few well-established markers for optimum concentrations. For a patient with a reliable diagnosis of bipolar disorder a common target for optimality would be blood concentration of 0.8-1 mM. Significantly higher concentrations will result in acute toxicity, while significantly lower will result in loss of effectiveness. However, this level has some side effects when sustained for years or decades, namely an increased risk of kidney damage and lowered thyroid activity. ^60^

At the other end of the dosage scale, epidemiological evidence is compelling that geographical variations in concentration of lithium in the drinking water are correlated with a variety of health and wellness markers, most notably and reliably with incidence of suicide. ^61,62,63 64 65 66^

Another important marker is provided by a study showing that over a four-year period a lithium level of .25-.4 mM of lithium (1/4 to 1/2 of the bipolar therapeutic dose) did not incur any renal damage^67^. This study suggests that clinical studies exploring low to medium-dose lithium could be undertaken with relatively minimal concerns for side effects.

One possible piece of low-hanging fruit for a clinical trial would be low- to medium-dose lithium for men undergoing active surveillance (AS) for advance of prostate cancer. From studies of AS outcomes, a large fraction of patients on AS ultimately require invasive treatment, as reviewed by Dall’Era et al^68^. When this need arises it typically comes after only a few years. Thus, a trial of lithium in this context would produce significant results in a short time and would be relatively inexpensive. One of us (EJ) conducted an informal one-person trial on himself after being diagnosed with prostate cancer in 2014, ingesting lithium supplements sufficient to bring his blood lithium to .3-.4mM while undergoing AS by Memorial Sloan Kettering Cancer Center. (MSK did not prescribe the lithium but agreed to include lithium level measurement in periodic blood tests.) In October 2017 EJ was told that there was no longer a need for AS. One case, important as it is to EJ, does not have statistical significance. We need clinical trials with significant numbers of people.

In ongoing work, we are constructing gene interaction networks built on a core based on those genes shared by the lithium-sensitive interactomes and the disease-specific networks; in other words, we will move from gene lists to gene networks. We will be happy to collaborate on further specific pathway or network analysis relevant to any of the cancers for which lithium may be a promising component of therapy.

## Author Contributions

The work was planned jointly in conversations between EJ and WG. WG did the computations and prepared the figures and tables. WG wrote the first draft of the Methods and Results sections. EJ wrote the first draft of the Introduction and Conclusions sections. Both authors shared in the final refinement of the manuscript.

